# Integrons, plasmids, and resistance genes in equine faecal bacteria

**DOI:** 10.1101/2022.09.12.507718

**Authors:** Scott W. Mitchell, Robert A. Moran, Liam D. H. Elbourne, Belinda Chapman, Michelle Bull, Gary Muscatello, Nicholas V. Coleman

**Author notes:** address correspondence to Nicholas Coleman.

## Abstract

Antimicrobial resistance in bacteria is a threat to both human and animal health. We aimed to understand the impact of domestication and antimicrobial treatment on the types and numbers of resistant bacteria, antibiotic resistance genes (ARGs), and class 1 integrons (C1I) in the equine gut microbiome. Antibiotic-resistant faecal bacteria were isolated from wild horses, healthy farm horses, and horses undergoing veterinary treatment, and isolates (9,083 colonies) were screened by PCR for C1I; these were found at frequencies of 9.8% (vet horses), 0.31% (farm horses), and 0.05% (wild horses). A collection of 71 unique C1I^+^ isolates (17 Actinobacteria and 54 Proteobacteria) was subjected to resistance profiling and genome sequencing. Farm horses yielded mostly C1I^+^ Actinobacteria (*Rhodococcus, Micrococcus, Microbacterium, Arthrobacter, Glutamibacter, Kocuria)*, while vet horses primarily gave C1I^+^ Proteobacteria (*Escherichia, Klebsiella, Enterobacter, Pantoea, Acinetobacter, Leclercia, Ochrobactrum)*; the vet isolates had more extensive resistance and stronger P_C_ promoters in the C1Is. All integrons in Actinobacteria were flanked by copies of IS*6100*, except in *Micrococcus*, where a novel IS*5* family element (IS*Mcte1*) was implicated in mobilization. In the Proteobacteria, C1I’s were predominantly associated with IS*26*, and also IS*1*, Tn*21*, Tn*1721*, Tn*512*, and a putative formaldehyde-resistance transposon (Tn*7489*). Several large C1I-containing plasmid contigs were retrieved; two of these (plasmid types Y and F) also had extensive sets of metal resistance genes, including a novel copper-resistance transposon (Tn*7519*). Both veterinary treatment and domestication increase the frequency of C1I’s in equine gut microflora, and each of these anthropogenic factors selects for a distinct group of integron-containing bacteria.

**IMPORTANCE:** There is increasing acknowledgement that a ‘One Health’ approach is required to tackle the growing problem of antimicrobial resistance. This requires that the issue is examined from not only the perspective of human medicine, but also includes consideration of the roles of antimicrobials in veterinary medicine and agriculture, and recognises the importance of other ecological compartments in the dissemination of ARGs and mobile genetic elements such as C1I. We have shown that domestication and veterinary treatment increase the frequency of occurrence of C1I’s in the equine gut microflora, and that in healthy farm horses, the C1I are unexpectedly found in Actinobacteria, while in horses receiving antimicrobial veterinary treatments, a taxonomic shift occurs, and the more typical integron-containing Proteobacteria are found. We identified several new mobile genetic elements (plasmids, IS and transposons) on genomic contigs from the integron-containing equine bacteria.

## INTRODUCTION

Antibiotic-resistant bacterial pathogens pose a serious threat to both human and animal health (1, 2). Tackling this problem effectively requires a One Health framework (3) that includes consideration of animals, plants, and environmental matrices (soil, sediments, sewage, water) as possible reservoirs and vectors for the resistant bacteria and their antibiotic resistance genes (ARGs) (4–6). A holistic approach is needed that includes not only human pathogens but also animal pathogens (7), the normal flora of humans and animals (5, 8, 9), and bacteria in the environment (10, 11). An improved understanding of the non-human reservoirs of resistant bacteria and ARGs is needed to develop mitigation strategies to limit their health impacts.

The problem of antibiotic resistance is partly due to the phenomenon of horizontal gene transfer (HGT) that allows bacteria to share advantageous traits and respond rapidly to environmental stresses (12). Mobile genetic elements (MGEs) including plasmids, phage, insertion sequences, transposons, gene cassettes and integrons are the major drivers of HGT in bacteria (13, 14); these elements act in a synergistic manner, leading to the development of complex mosaic elements (15, 16). Integrons in particular enable bacteria to capture, shuffle, and express ARGs packaged as modular gene cassettes (17, 18). The class 1 integron (C1I) is the most widely disseminated type (19), and is capable of assembling large ARG arrays conferring extensive antibiotic resistance (20). New genetic contexts and bacterial hosts for C1Is are constantly being discovered (21), and thus these elements pose an evolving and ongoing health threat.

The role of animals as reservoirs and vectors for the transmission of C1Is and their associated ARGs is well established. Prior studies have focused on food animals (e.g. pigs, poultry and cattle (22, 23) and have highlighted problems with intensive factory farming practices which lead to the spread of C1Is and ARGs into the food chain (24) or the environment (25). It is also clear that companion animals and wild animals can carry C1Is and ARGs, with previous studies highlighting their presence in dogs (26), cats (27), rabbits (28), horses (29), bats (30), seals (31), fish (32) and birds (33), among others. The overall picture is grim, and the global dissemination of C1Is and other ‘xenogenetic DNAs’ (34) into most ecosystems and animal species has led to these being considered as a new kind of persistent pollutant (35, 36).

Horses are a relatively understudied animal system with respect to the problem of antibiotic resistance (37), although like most animals, they are known to be a reservoir of resistant bacteria, ARGs and C1Is (29, 38–41). Many of the same classes of antibiotics are used in both human and equine medicine (37), and thus it might be hypothesised that these drugs may select for bacteria, MGEs or ARGs that later contribute to problems for human health. Rigorous evidence to support this hypothesis is lacking, and an outstanding question relates to the relative roles of antimicrobial exposure versus general human contact (i.e. domestication) in selecting for particular types of resistance phenotypes and genotypes. Previous attempts to address this question in horses and other animals have been limited by small sample sizes, a lack of controls (i.e. wild animals of the same species), a paucity of genomic and phenotypic data, and/or the targeting of specific species (usually *Escherichia coli*) at the expense of other microflora that may be present.

In this study, we aimed to elucidate the factors that impact the types and numbers of antibiotic-resistant bacteria, C1Is, and ARGs in equine faecal samples. To address the limitations described above, we sampled three different cohorts at differing levels of anthropogenic impact (wild horses, healthy farm horses, and sick horses undergoing veterinary treatment) using general-purpose rather than taxon-selective media for isolation, and including extensive phenotypic and genomic analyses on the resultant C1I-containing isolates.

## RESULTS

### Antibiotic resistance in equine faecal bacteria

Faecal samples were collected from three horse cohorts representing different levels of anthropogenic impact; these included 11 wild horses from a national park, 11 healthy farm horses, and 11 sick horses undergoing veterinary treatment. The collection locations are shown in Supplementary Figure 1. Dilutions of equine faecal samples plated onto R2A agar containing different antibiotics yielded numerous and diverse resistant colonies. Viable counts of resistant bacteria were significantly higher (p<0.005) in samples from the vet and farm cohorts compared to the wild cohort for all antibiotics except streptomycin (Figure 1A, see Supplementary Table 1 for p values). The data were organised into an ‘increased human exposure’ group (farm and vet cohorts, n=176) and a ‘decreased human exposure’ group (wild cohort, n=88) to facilitate further statistical analysis. The numbers of resistant bacteria averaged across all antibiotics tested were higher in the ‘increased exposure’ group (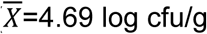, SD=1.26) relative to the ‘decreased exposure’ group (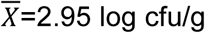, SD=1.09), at a significance level of p <0.001.

**Figure 1.**
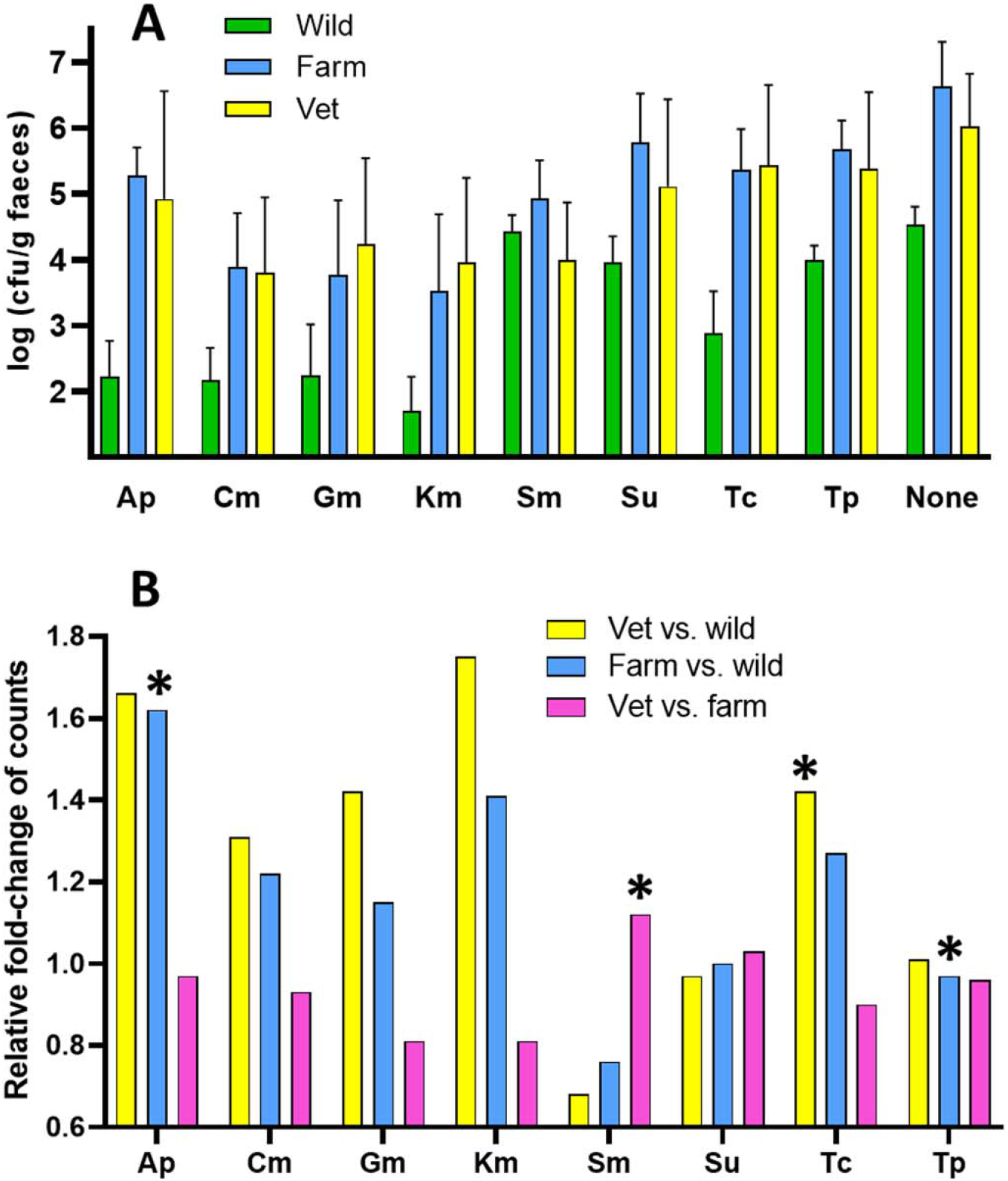
Comparison of resistant bacteria in different equine cohorts. **A**. Numbers of colonies on different antibiotic agars. **B**. Relative fold-changes after normalisation to counts on no-antibiotic agar plates. Columns with error bars indicate the mean and SD, respectively (n = 11). Asterisks indicate significant differences based on normalised error thresholds (Supplementary Figure 1).

A complicating factor in the above analyses is that viable counts on plates with no antibiotics added were higher for the farm and vet horse samples compared to the wild horse samples. Therefore, an alternative analysis was done by calculating the fold-changes in viable counts relative to the fold-changes in the no-antibiotic controls (Figure 1B.). At face value, this analysis showed higher fold-changes in resistant bacteria in the vet vs. wild and farm vs. wild comparisons for all antibiotics except streptomycin, sulfamethoxazole, and trimethoprim, but a more rigorous normalised error analysis (Supplementary Figure 1) indicated that only a handful of these changes were statistically significant (Ap^R^ in farm vs. wild, Sm^R^ in vet vs. farm, Tc^R^ in vet vs. wild, and Tp^R^ in farm vs. wild). Taken together, it appears that human association and veterinary treatment do increase the frequency of at least some kinds of antibiotic-resistant bacteria in equine faecal samples, but the effect is not strong and is not uniform for all antibiotic types.

We next searched for correlations between the veterinary use of particular antibiotics and the abundance of the corresponding resistant bacteria (Supplementary Table 2). The only antibiotic that gave a significant effect here (p<0.05) was Gm. The counts of Gm^R^ bacteria in samples from horses treated with gentamicin within 1 month of sample collection (n=5, 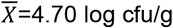, SD=1.30) were significantly increased (p=0.011) compared with those from untreated horses (n=28, 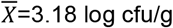, SD=1.27). This selective effect seemed to fade over time, since reanalysis of the data based on horses treated with Gm within 4 months of sampling did not reveal the same trend.

### Quantification and characterisation of class 1 integron-containing isolates

The frequencies of detection of C1Is by PCR in colonies arising on antibiotic-containing R2A agar plates were dramatically different between cohorts; wild horses yielded only 2/2484 positives (0.081%), while healthy farm horses yielded 18/2784 positives (0.65%) and horses undergoing veterinary treatment yielded 345/3815 positives (9.04%). The C1I-containing bacteria were only found on sulfamethoxazole plates in the samples from wild horses, but they were most frequent on tetracycline plates in samples from the farm horses, and on gentamicin plates in the samples from vet horses (see Supplementary Table 3 for details). After dereplication by BOX PCR, the CI1-containing strain collection consisted of 1 isolate from the wild cohort, 13 from the farm cohort, and 57 from the vet cohort (71 in total). The C1I-containing bacteria were identified by partial 16S rDNA sequencing and/or draft genome sequencing as representing 8 genera of Proteobacteria (*Escherichia, Klebsiella, Enterobacter, Pantoea, Leclercia, Atlantibacter, Acinetobacter, Ochrobactrum)* and 6 genera of Actinobacteria (*Rhodococcus, Micrococcus, Microbacterium, Arthrobacter, Glutamibacter, Kocuria*). The majority of isolates from the farm cohort (11/13) were Actinobacteria, while the majority of isolates from the vet cohort (51/57) were Proteobacteria (Table 1, Supplementary Table 4).

**Table 1.**
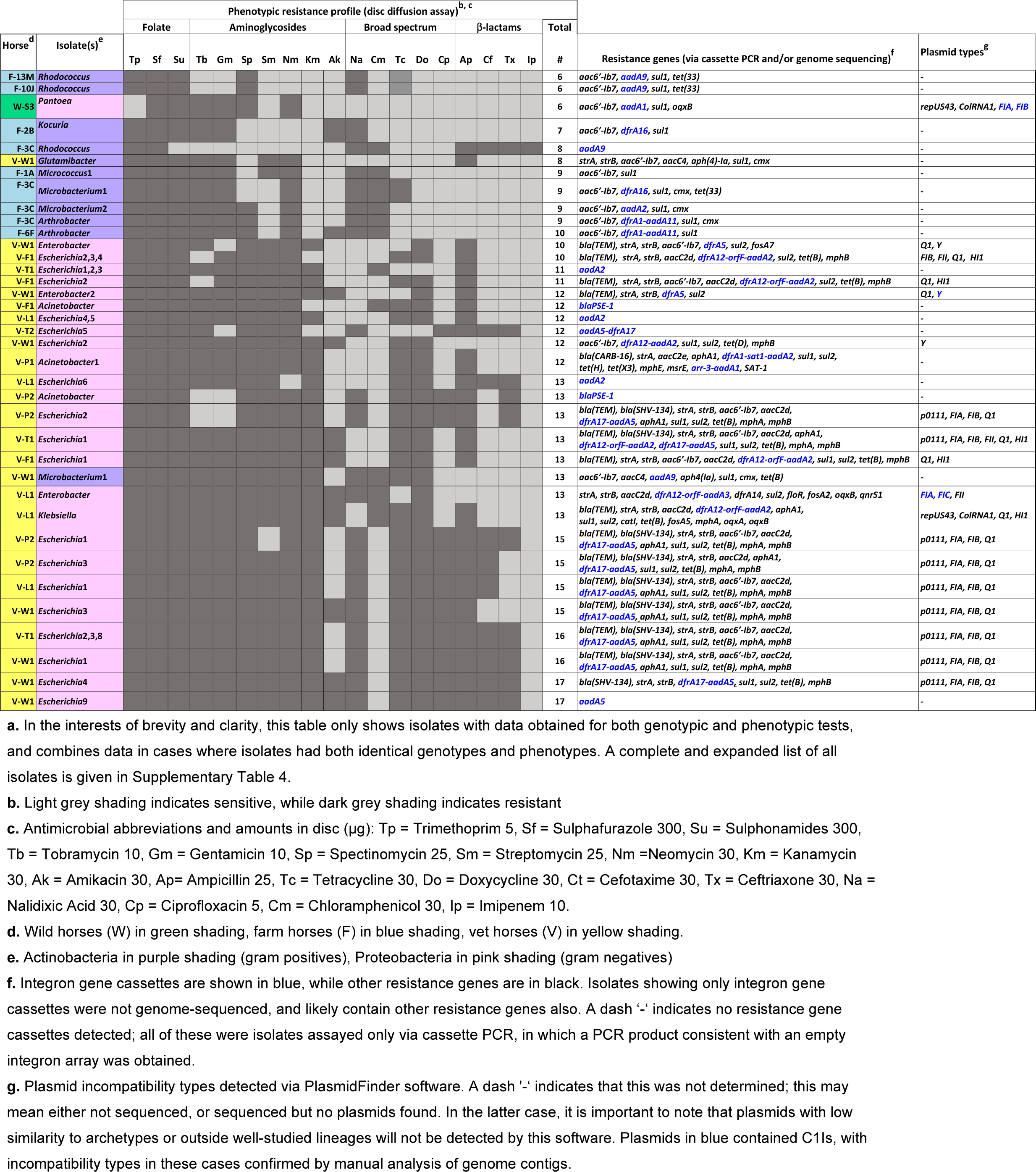
Antimicrobial resistance phenotypes and genotypes of C1I-containing equine isolates^a^

The C1I^+^ isolates were tested for resistance against a panel of 19 antibiotics (Figure 2, Table 1). Isolates from vet horses tended to have more resistances (12.7 ± 2.7) than those from farm horses (7.6 ± 2.0), while the single C1I^+^ isolate from a wild horse had 6 detected resistances. Proteobacteria tended to have more resistances (12.9 ± 2.7) than Actinobacteria (7.9 ± 2.2); the fact that these numbers are nearly identical to those based on cohort simply reflects that most C1I^+^ vet isolates were Proteobacteria and most C1I^+^ farm isolates were Actinobacteria. Resistance to trimethoprim and sulfonamides was very common in all isolates (>90% of isolates resistant), and resistance to aminoglycosides (except amikacin) was common in the vet cohort isolates (>70% of isolates resistant). Resistance to broad-spectrum antibiotics like chloramphenicol and tetracycline was seen in many isolates from both farm and vet cohorts.

**Figure 2.**
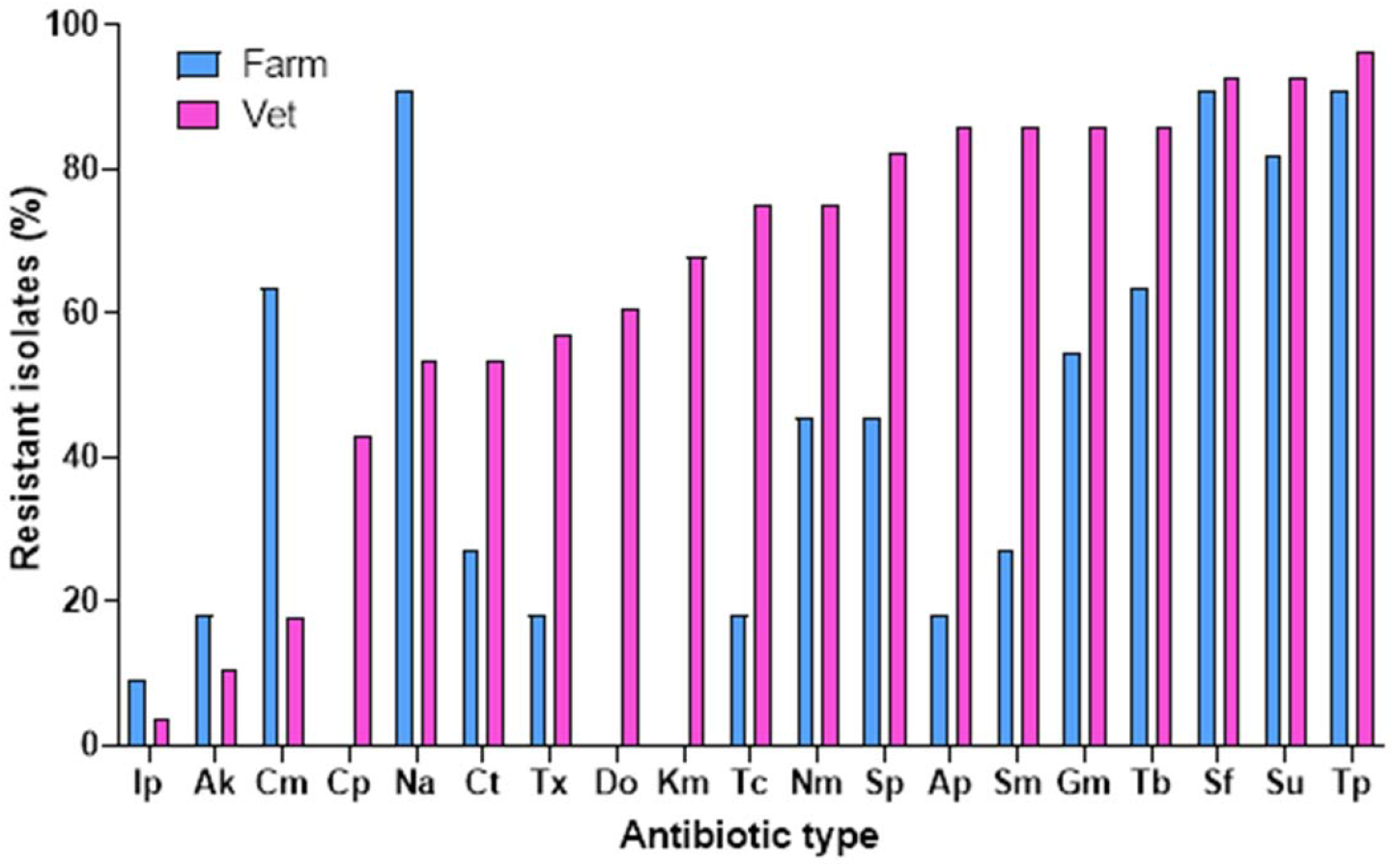
Antibiotic resistance of C1I-containing isolates in farm vs. vet cohorts Bars indicate the percentage of C1I-containing isolates from each cohort that were resistant to each individual antibiotic. The single wild cohort isolate is not included in this analysis; this isolate was resistant to Su, Sf, Tb, Gm, Sm, and Nm.

The integron gene cassette content of all C1I+ isolates was assessed by PCR, using a forward primer in the 5’ conserved segment (5’ CS) and a reverse primer in the 3’ conserved segment (3’CS) (58) (Table 1, Supplementary Table 4); 49 isolates contained gene cassettes, 3 isolates had no cassettes in the integron, and 19 isolates gave no PCR product. The latter situation indicates most likely that deletions involving the *attI1* site and/or *qacEΔ* gene had occurred. Streptomycin resistance (*aadA*) and trimethoprim resistance (*dfrA*) gene cassettes were by far the most frequently encountered, as seen in many other studies, and a variety of specific sequence variants of these were seen. The only other genes encountered in C1I cassette arrays were *orfF* (unknown function), *bla*_PSE-1_ (extended-spectrum beta-lactamase), and *arr3* (rifampicin resistance).

Draft genome sequencing of a subset of the C1I+ isolates (33 in total) confirmed the findings from cassette PCRs and revealed many further resistance genes (Table 1, Supplementary Tables 4 and 5), including some cassettes (*dfrA1-sat1-aadA2*) associated with a class 2 integron that were not retrieved by the class 1 integron-specific PCR. Very extensive sets of resistance genes (up to 16) were detected in draft genomes from many of the vet cohort isolates. There was approximate agreement between the numbers of phenotypic resistances detected and numbers of resistance genes detected in genomes, with the farm cohort isolates yielding fewer resistance genes (3.9 ± 0.9) than the vet cohort isolates (10.5 ± 3.8) (note that these calculations only include genome-sequenced isolates). Several resistance gene profiles were identical across multiple independent isolates (two *Rhodococcus* isolates from farm horses 10J and 13M, three *Escherichia* isolates from vet horses T1 and P2, six *Escherichia* isolates from vet horses L1, P2, T1, W1; Supplementary Table 5); this strongly suggests movement of resistant bacterial clones or MGEs between individual animals.

### Analysis of integron-containing genome contigs from Proteobacteria

#### Escherichia

Draft genome sequences were obtained from 16 distinct C1I^+^ *Escherichia* isolates from five vet horses. Identical 4539 bp CI1-containing contigs (V-L1-*Escherichia*1; V-P2-*Escherichia*1,2,3; V-T1-*Escherichia*2,3,8; V-W1-*Escherichia*1,3,4) were found in 10 isolates; these contained *dfrA17* and *aadA5* cassettes, a 5’ conserved region (5’-CS) up to the IRi repeat, and a 3’ conserved region (3’-CS) with *qacE*Δ, *sul1*, and *orf5* (Figure 3a). Partial copies of IS*26* are at each end of the contig and thus this C1I is likely to be the passenger segment in an IS*26* pseudo-compound transposon (59). Five *Escherichia* contigs (V-F1-*Escherichia*1,2,3,4; V-T1-*Escherichia*1.1) had a different sequence (3660 bp), this was a C1I with *dfrA12, orfF*, and *aadA2*, and a shorter 3’ CS, truncated by IS*1* (Figure 3b). A single *Escherichia* gave a third type of contig (V-W1-*Escherichia*2; 7617 bp); this contained *dfrA12, orfF*, and *aadA2*, and was embedded in Tn*21*, which had been truncated by IS*1* at the 5’ end and IS*26* at the 3’ end (Figure 3c).

**Figure 3.**
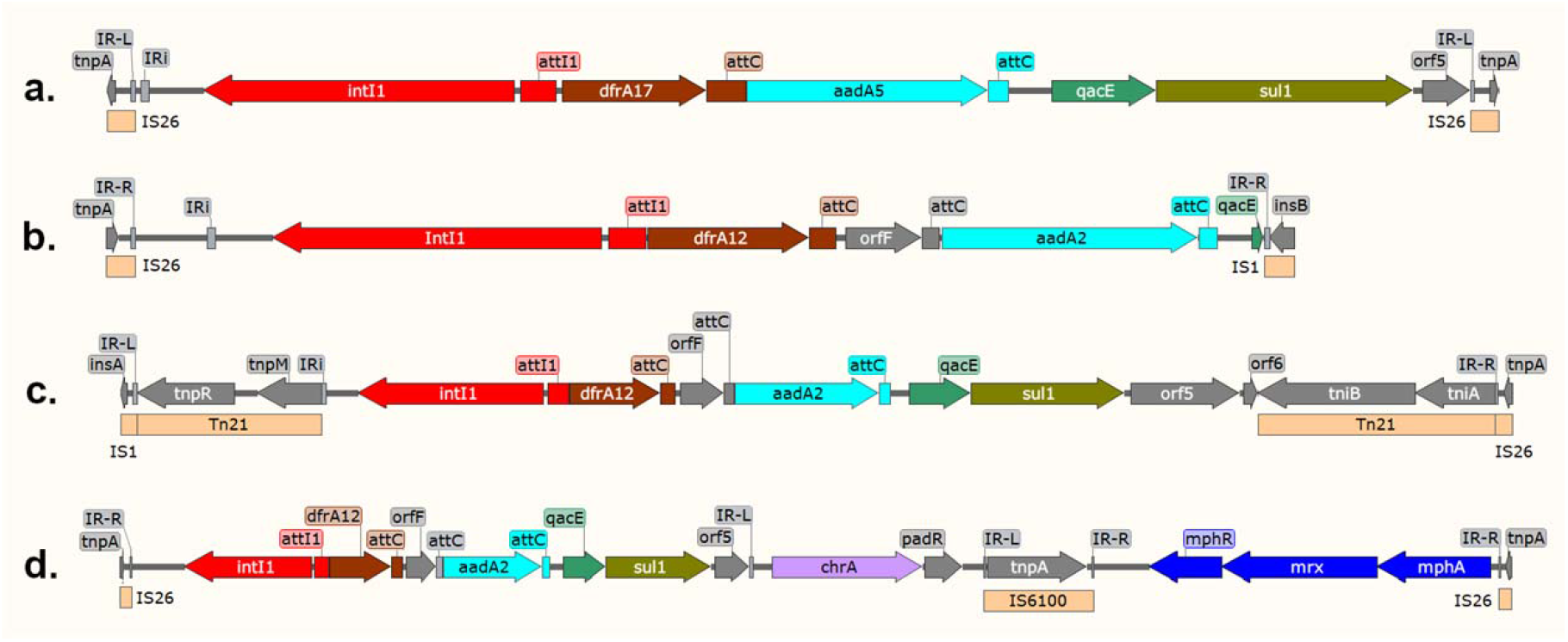
Integron-containing contigs from *Escherichia* and *Klebsiella* isolates (a) V-P2-*Escherichia*1 (b) V-F1-*Escherichia*1 (c) V-W1-*Escherichia*2 (d) V-L1-*Klebsiella*

#### Klebsiella

A single C1I^+^ *Klebsiella* yielded an 11,088 bp contig (V-L1-*Klebsiella*; Figure 3d). The first part (2956 bp) was the same as V-F1-*Ecoli*1, but this was not truncated by IS1, and continued into a chromate resistance gene (*chrA*), IS*6100*, then a macrolide resistance cluster (*mphR-mrx-mphA*), then the contig ends at IS*26*. As above, the whole contig is likely to be part of an IS*26*-based transposon. This same region was found in approximately 150 other *Enterobacteriaceae* plasmids and genomic islands via blastn analysis, and has been previously well-characterised in human isolates of *E*.*coli* (60) and *Klebsiella pneumoniae* (61) and in porcine *Aeromonas hydrophila* (62).

### Enterobacter

One C1I^+^ *Enterobacter* isolate from horse V-L1 yielded a 50,408 bp contig (V-L1-*Enterobacter*; Figure 4a) that contains a fragment of a C1I (*attI1*-*dfrA12*-*orfF*-*aadA2*) truncated at the left end by the contig break, and at the right end by an IS*1* insertion. This contig is clearly part of a plasmid, and a large region (43 kb) is near-identical to two other thus-far uncharacterised *Enterobacter* plasmids (unpublished, MZ156800, CP019890). The plasmid has FIA and FII replication functions. The FIA-type *repE* gene and the adjacent *oriV* / *incC* regions are very similar to the F plasmid of *E*.*coli* (96% inferred amino acid (aa) identity in RepE) while the FII RepA protein has 64% inferred aa identity to that of *E*.*coli* enterotoxin plasmid EntP307. The contig contains multiple stability and partitioning genes (*parA, parB, stbA*), a toxin/antitoxin system (*higA-higB*) and one putative conjugation gene (*traX*).

**Figure 4.**
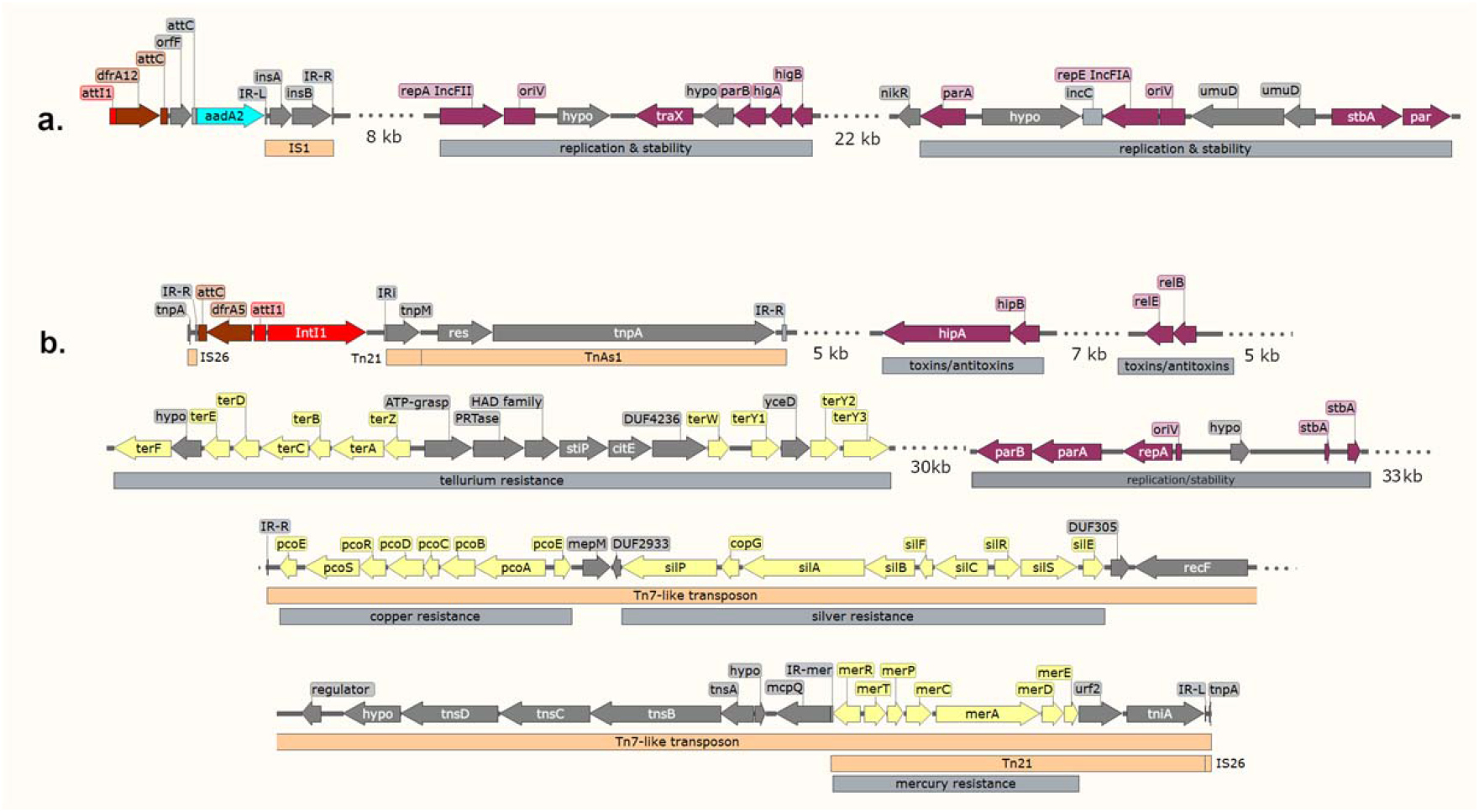
Features of integron-containing contigs from *Enterobacter* isolates. (a) V-L1-*Enterobacter* (b) V-W1-*Enterobacter2*

Two distinct *Enterobacter* strains from horse V-W1 yielded contigs V-W1-*Enterobacter*1 (88,930 bp) and V-W1-*Enterobacter*2 (149,407 bp, Figure 4b); these were identical over the region shared, so just V-W1-*Enterobacter*2 is discussed further here. This contig shared some common regions with other *Enterobacteriaceae* plasmids, but the configuration of elements in this whole structure was unique based on blastn comparisons to the GenBank database. The C1I contained *dfrA5*, and was associated with a recombinant Tn*21*:Tn*1721* element; this configuration has been described previously (63). The 3’CS is truncated by IS*26*, which breaks the contig on the left. A *repA* gene encodes a protein with 91% inferred aa identity to that of P1, meaning this contig is part of a Y-type phage-plasmid. Partitioning genes *parA* and *parB* are present, along with two toxin/antitoxin systems (*hipB-hipA* and *relB-relE*), and two *traG* homologs (*virD4*). Two putative tellurium resistance gene clusters (*terZABCDEF, terWY1Y2Y3*) are present from 25-42 kb, and a large (21 kb) silver and copper resistance region is present between 110-132 kb (*silESRCFBAP* and *pcoEABCDRSE*). The copper and silver resistance genes are part of a Tn*7*-like transposon. Similar transposons are seen in many enterobacterial plasmids e.g R478 (64, 65). A section of Tn*21* is embedded within the Tn*7*-like transposon including the mercury resistance genes (*merRTPCAD*). It is not clear from the draft genome sequence whether this is the same copy of Tn*21* also seen at the beginning of the contig. The Tn*21* is truncated by IS*26*, which ends the contig on the right.

### Acinetobacter

A single *Acinetobacter* isolate yielded three integron-containing contigs (V-P1-*Acinetobacter*1.1, 1.2, 1.3; Figure 5). Contig V-P1-*Acinetobacter*1.1 contains a class 2 integron (*dfrA1*-*sat2*-*aadA1*) inside Tn*7*, then *sul2*, then *tet(X3)* flanked by two ISCR2 elements. Tet(X3) confers resistance to tigecycline, an important last-resort antibiotic for *Acinetobacter* infections, so this finding is of concern. Adjacent to *tet(X3)* is *parA*, with 82% inferred aa identity to ParA of RK2/RP4; this suggests the contig is part of an P-type plasmid. The leftmost 6.5 kb of contig V-P1-*Acinetobacter*1.1 is found in approximately 20 other *Acinetobacter* genomes, but its position relative to a class 2 integron appears to be unique here. V-P1-*Acinetobacter*1.2 and V-P1-*Acinetobacter*1.3 are likely to be consecutive genome regions, and contain a derivative of Tn*21* (∼14 kb), which carries a C1I with *arr3* and *aadA1* cassettes, and several insertion sequences; IS*UnCu1* (inserted in the *aadA1 attC* site), and IS*Pa110* (inserted into IS*1321*) (66, 67). This particular Tn*21* derivative is also seen in three other sequences in GenBank (CP026932, CP038791, CP027355), from human and bovine *E*.*coli* isolates. Outside the Tn*21* sequence are IS*Alw33*, an IS*5-*family element similar to IS*Abe15*, and a fragment of IS*Aba1*.

**Figure 5.**
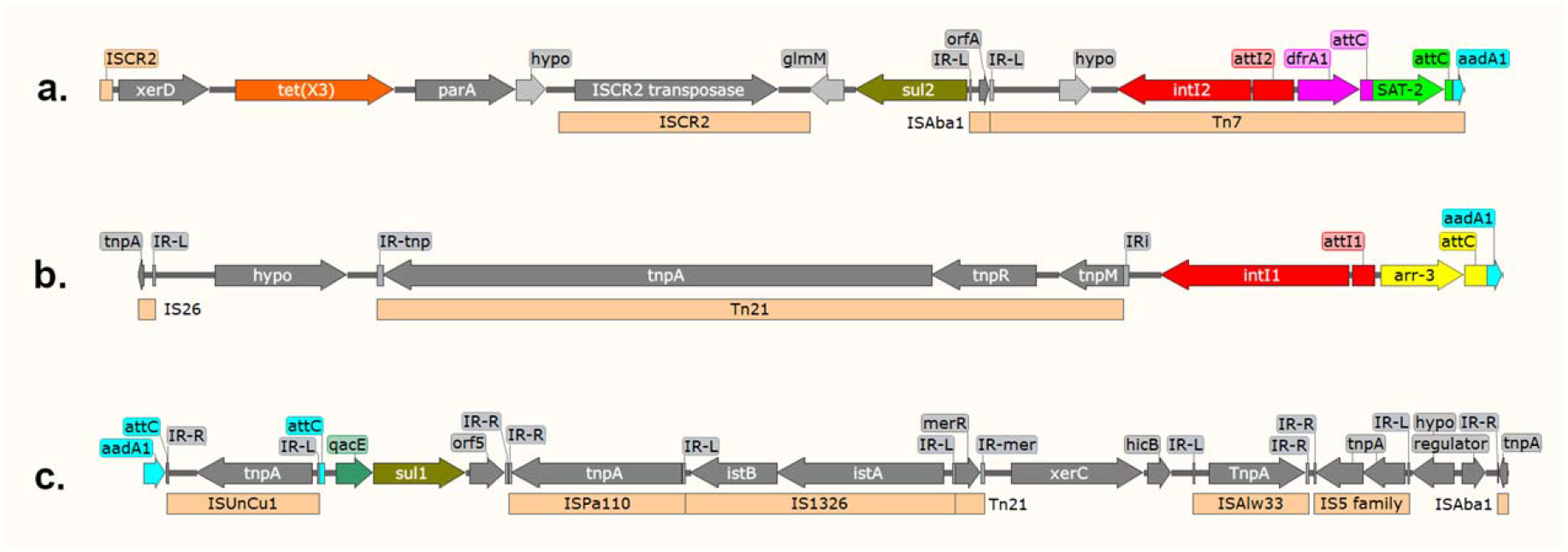
Integron-containing contigs from an *Acinetobacter* sp. a. V-P1-*Acinetobacter*1.1 b. V-P1-*Acinetobacter*1.2 c. V-P1-*Acinetobacter*1.3

### Pantoea

A single large C1I^+^ contig (W-S3-Pantoea; 184 kb) was obtained from an isolate from the wild horse cohort (Figure 6). This is a section of an FIA/FIB plasmid, with RepE and RepB exhibiting 29% and 76% aa identity, respectively, to those of F plasmids. The replication origins (*oriS* and *oriV*) are similar also to F plasmids (68); the *oriV* has 5 repeats upstream, and 8 repeats downstream of *repB*, with consensus ATAAGCTATAG (almost identical to the ATAAGCTGTAG in plasmid F). A putative *oriS* is upstream of *repE*, consisting of a 100-bp AT-rich region with 4 repeats (consensus TTTAAAGCTT). Genes for partitioning functions *sopA, stbA, stbB* are present, similar to F plasmids. A single putative conjugation gene is present, encoding TraT with 72% aa inferred identity to that of R100.

**Figure 6.**
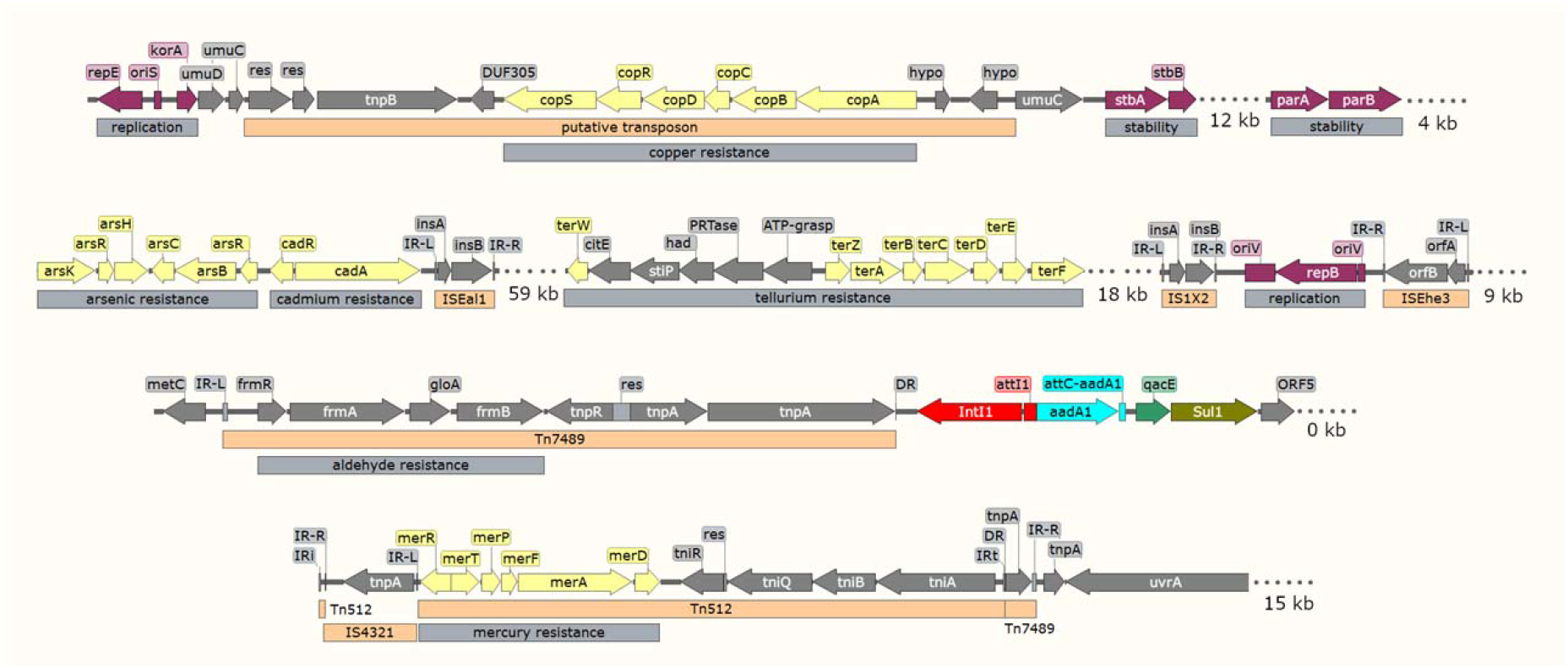
Features of integron containing contig W-S3-*Pantoea*

The *Pantoea* C1I is associated with a Tn*512*-like element, inserted inside a Tn3 family transposon that has been deposited in the Transposon Registry (69) as Tn7489. The Tn*512* and C1I appear to have moved as a composite unit (13,637 bp), using the IRi of the integron as the left end, and the IRt of Tn*512* as the right end; evidence for this comes from 5 bp direct repeats (AACGC) adjacent to these ends, and four other database examples can be found of similar events. Tn*7489* encodes a DDE transposase and a serine resolvase with 90% and 96% predicted aa identities, respectively, to those of Tn*Ec1* (70), and its terminal 47 bp inverted repeats are near-identical to Tn*Ec1*. Unlike Tn*Ec1*, Tn*7489* is a genuine transposon, in the sense that it has cargo genes (*frmR, frmA, gloA, frmB*). The *tnpA* of Tn*7489* is a pseudogene, and contains an internal stop codon in addition to being interrupted by the Tn512-like transposon.

Structures similar to Tn*7489* (>97% coverage and 90-96% nucleotide identity) exist in 106 other *Enterobacteriaceae* plasmids, including well-studied elements such as R471 (the IncL archetype) and Rts1 (the IncT archetype). This type of Tn has previously been formally named Tn*6901* (71) and more recently renamed Tn*6399* (72) but their predicted phenotype was not clear. One recent report (73) detected the *frmR-frmA-gloA-frmB* gene cluster in a small *Aeromonas* plasmid pAsaXII, and confirmed that these genes confer formaldehyde resistance, although in that case there was no Tn structure surrounding them. That advance now enables us to confidently predict that Tn*6901* / Tn*6399* / Tn*7489* are formaldehyde-resistance elements.

A striking feature of the *Pantoea* contig is multiple metal-resistance regions, including tellurite (*terWZABCDEF*), copper (*copABCDRS*), arsenic (*arsKRHCBR*) and cadmium (*cadRA*) resistance. The arsenic and cadmium resistance region (8 kb) is similar to those found in many Gram negative plasmids. The tellurite resistance region (12 kb) has only three close matches in GenBank, from *Pantoea* and *Acinetobacter* plasmids, but these are either unpublished (CP083449, CP060163) or reported but not well characterised (CP010325) (74). The copper resistance region (6 kb) is inserted precisely within *umuC* and associated with a Tn*554-*like transposase; this appears to be a novel copper resistance transposon (note that Tn*554* family elements lack IRs or DRs, making it difficult to define the ends); we have deposited this in the Transposon Registry as Tn*7519*. The other main phenotypes predicted to be encoded by this *Pantoea* plasmid are the metabolism and transport of sugars, since homologs of *srlL, glmS, xylE, gdhI, frlRDBA, amyA, budRAB, gcd, citE, pfkB, galM, lamB, sacC, fruR* are present (see GenBank OP296273; not shown in Figure 8).

**Figure 7.**
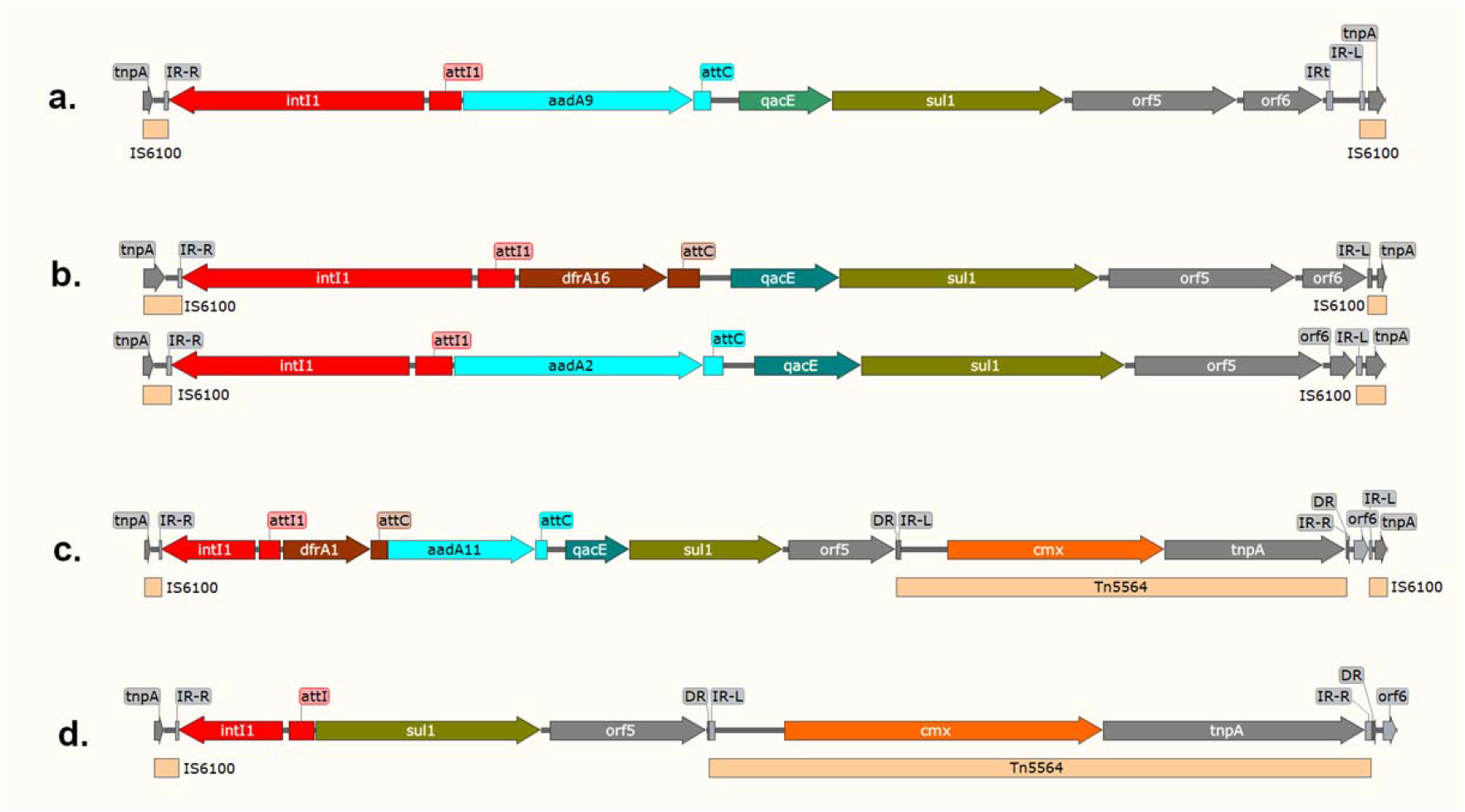
Integron-containing contigs from *Rhodococcus, Microbacterium, Arthrobacter* and *Glutamibacter* a. F-10J-*Rhodococcus* and F-13M-*Rhodococcus* b. F3C-*Microbacterium*1 and F-3C-*Microbacterium*2 c. F-3C-*Arthrobacter* and F-3C-*Arthrobacter* and V-T1-*Arthrobacter* d. V-W1-*Glutamibacter*

**Figure 8.**
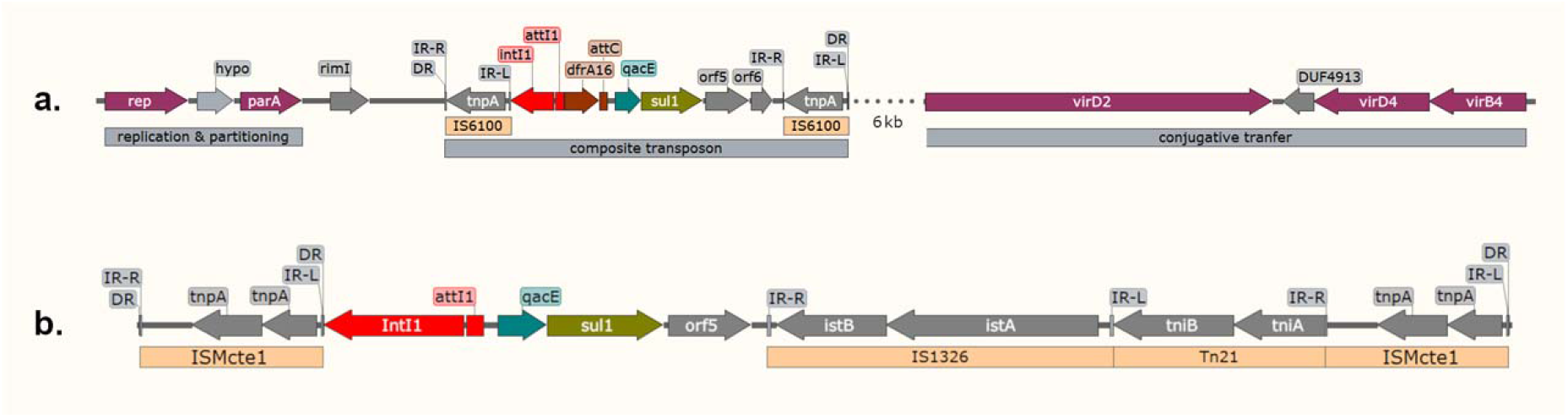
Features of integron-containing contigs from *Kocuria* and *Micrococcus*. a. F-2B-*Kocuria* b. F-1A-*Micrococcus*

### Analysis of integron-containing contigs in equine Actinobacteria

Eleven C1I-containing contigs were obtained from eleven distinct actinobacterial isolates including *Arthrobacter, Glutamibacter, Micrococcus, Rhodococcus, Microbacterium*, and *Kocuria* spp (Figure 7, Figure 8). In all these cases except for *Micrococcus*, the C1I are flanked by copies of IS*6100* and appear to be part of composite transposons. Similar elements have been reported previously in *Corynebacterium* (75) and *Rothia* (76). Comparison of the integron-IS*6100* junctions (Figure 9) indicates that many different IS6100-mediated events have occurred to generate these different contigs, although in three cases, different contigs feature the same integron-IS*6100* junction at one end.

**Figure 9.**
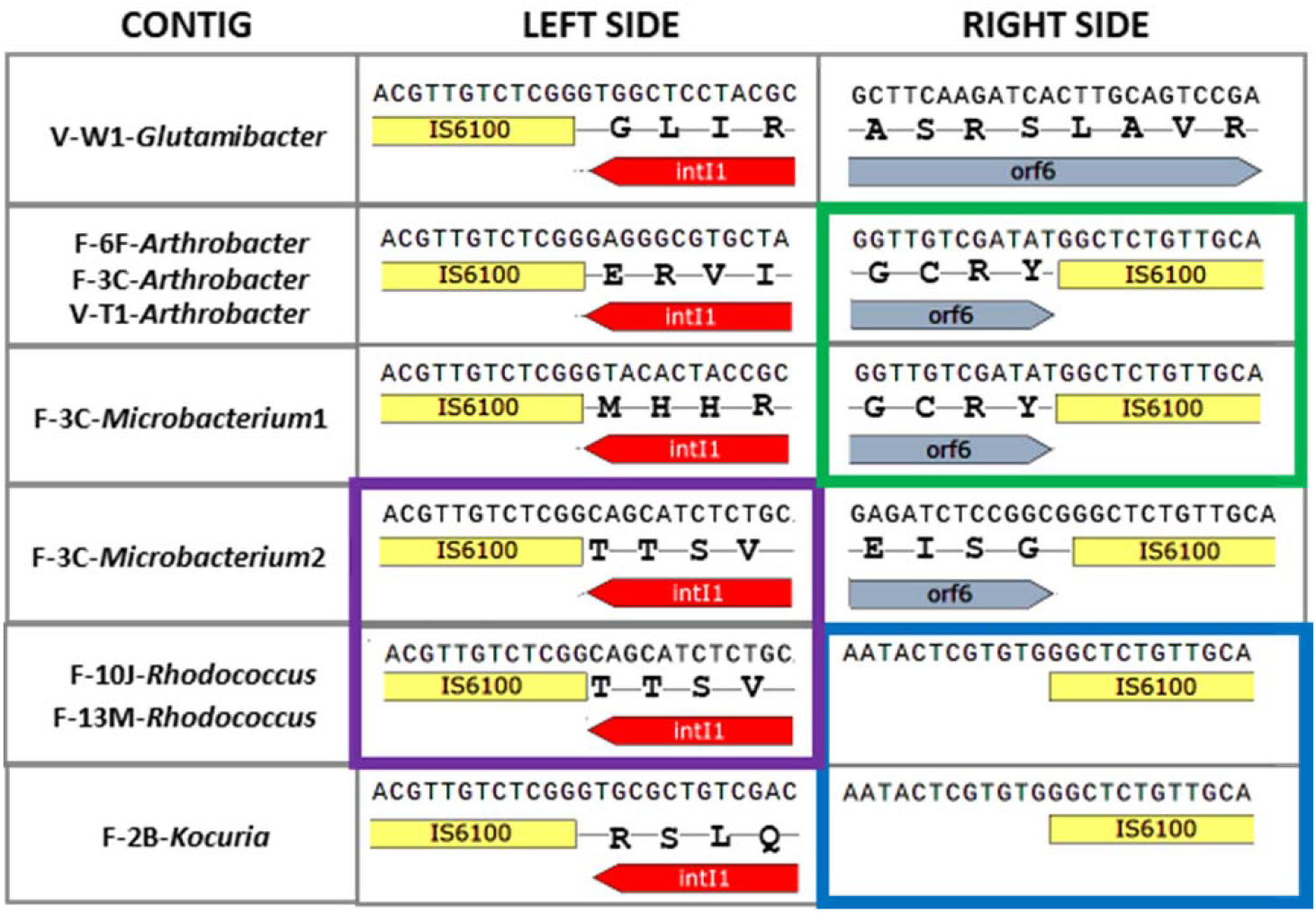
Integron-IS*6100* junctions in *Actinobacteria*. Ends that match are linked by the coloured boxes (green / purple / blue)

The contigs from Actinobacteria contain *dfrA16, aadA2* or *aadA9* cassettes, and all have an intact 3’ CS (*qacEΔ, sul1, orf5*). Evidence for the spread of specific C1I’s between horses and between cohorts was found in these Actinobacteria; identical *aadA9*-containing contigs (4504 bp) were obtained from *Rhodococcus* isolates from farm horses F-10J and F-13M, and identical *dfrA1-aadA11* containing contigs (6733 bp) were obtained from *Arthrobacter* isolates from horses F-3C, F-6F and V-T1. The three identical *Arthrobacter* contigs contain Tn*5564* (*cmx*, Cm^R^) inserted in *orf5*; evidence for transposition is seen by the flanking 6 bp DR’s CCTAGA. A contig from *Glutamibacter* (V-W1) was similar to the *Arthrobacter* sequences but had a 1912 bp internal deletion, a different junction between IS*6100* and *intI1*, and a slightly different Tn*5564* sequence (5 nucleotide differences over 2438 bp).

The F-2B-*Kocuria* contig (Figure 8a) includes complete sequences of both IS6100 elements, enabling identification of 8-bp flanking DRs at the outer edges, thus providing evidence for movement of this element as a composite transposon. Plasmid genes for replication, partitioning and transfer are nearby; the closest homologs are RepA and ParA in the *Rhocococcus equi* virulence plasmid pVAPN1572 (27% and 21% inferred aa identity, respectively) and TraA/VirD2 of *Rhodococcus erythropolis* plasmid pREA400 (27% inferred aa identity). More distant homologs of all these genes also occur in *Corynebacterium* spp. resistance plasmids (pNG2, pTET3) (75, 77).

The F-1A-*Micrococcus* contig (Figure 8b) contains a C1I with no cassettes, attached to IS*1326* and Tn*21*. These parts together have 99% nucleotide identity to In0 from *Pseudomonas* pVS1.The CI1 and Tn*21* parts are flanked by two identical copies of a novel IS*5* family element, which has been deposited in the IS Finder database, and named IS*Mcte1*. This element has identical 16 bp IR’s, generates a 4 bp DR (CTAG) upon transposition, and has a DDE transposase encoded by three overlapping ORFs. The closest characterised match to IS*Mcte1* was IS*Bli8*, with 52% aa identity in the transposase. The putative IS*Mcte1*-based transposon is chromosomal; evidence for this comes from the lack of any obvious plasmid genes on this large contig (345 kb), and the presence of several essential genes e.g. ribosomal proteins (*rplY*), cell division functions (*ftsL*), and tRNA’s (Asn, Gln, Leu, Arg). Figure 8 only shows a small region of interest from the *Micrococcus* C1I-containing contig; for more information, see GenBank OP296261.

### Pc promoter variants in the equine integron collection

The Pc promoter drives expression of integrated cassettes in integrons. There are five distinct Pc sequence variants in the C1I^+^ equine isolates (Figure 10). There was a trend for Proteobacteria to have the strong Pc variants and Actinobacteria to have the weaker Pc variants. Two sequence variants of PcW are present, one has an A→C at the -27 position; it is not clear whether this impacts function. Pc is located within the *intI1* gene, and so there is a tension between evolving towards stronger Pc variants versus evolving towards more active IntI1 integrases (78). This may explain why a diversity of Pc promoters persist in C1I’s in this strain collection and in the wild.

**Figure 10.**
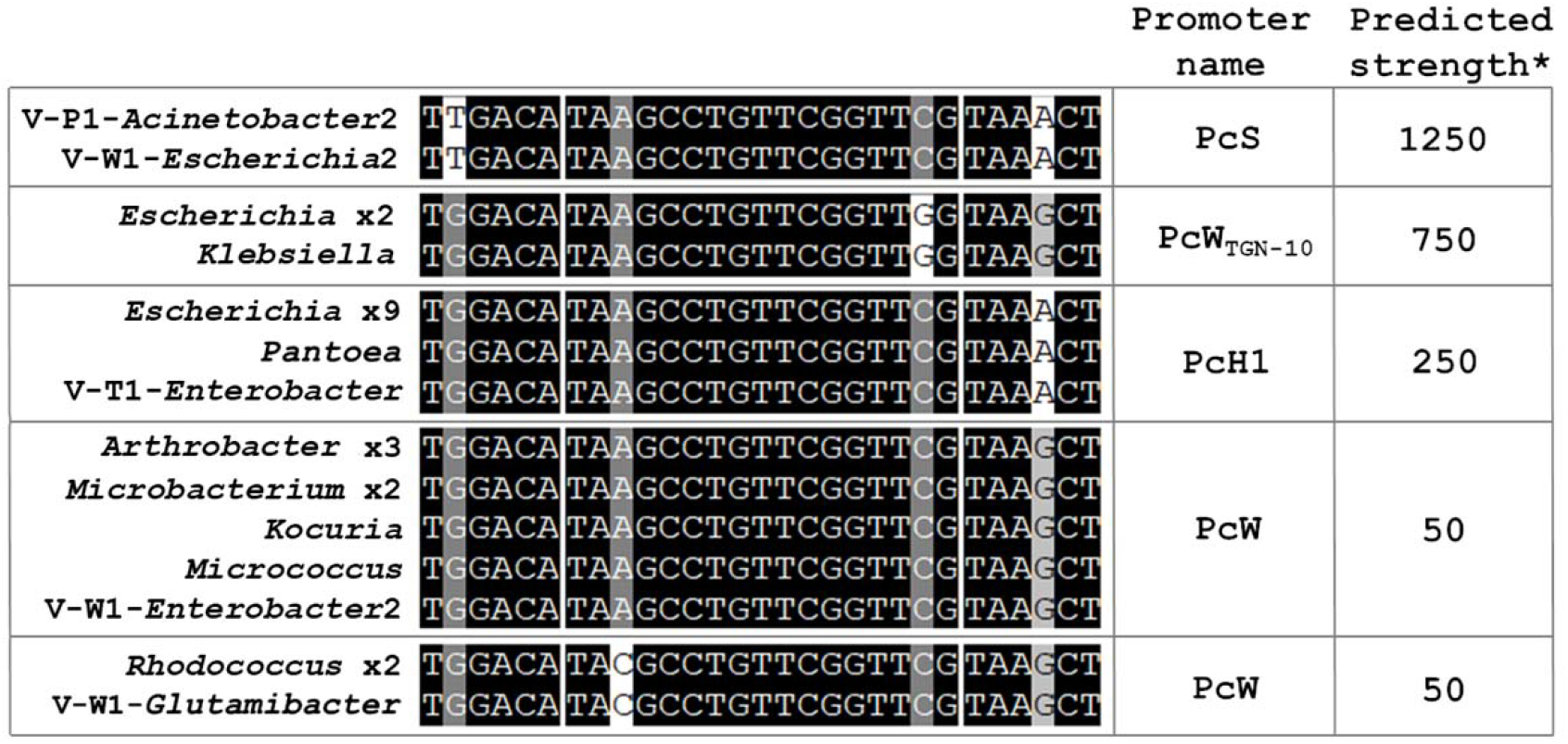
Pc promoter variants in the equine C1I collection. Data from 27 contigs are included. Multiple contigs are shown as e.g. *Escherichia* x9. *Predicted strength from Ploy et al, 2010 (Miller units from Pc-lacZ fusions).

## DISCUSSION

Our study confirms the importance of C1Is in the persistence and spread of ARGs in equines and highlights the role of equine gut microflora as a reservoir for many different MGEs and ARGs that may impact on animal and human health. Class 1 integrons are known to be an excellent marker for anthropogenic impacts (35) and a strong effect of human contact on C1I abundance has been seen in studies of other animals (28, 30, 79–82). We have now extended these findings to the case of equines, where C1Is were seen to be 8-fold and 112-fold more abundant in farm and vet horses, respectively, compared to wild horses. The situation with resistance phenotypes was less clear-cut, with only Tc^R^, Ap^R^ and Tp^R^ displaying statistically significant increases in the farm or vet horses versus wild horses, after correction for differences in overall viable counts between cohorts. In the case of the vet horses, increases in C1I frequency and resistance could be due to direct selective pressure from antibiotic treatment (83), while for both vet and farm horses, other factors may include contamination of soil or feed with antibiotic residues or ARGs (84, 85), acquisition directly from humans (86), or transmission from other farm animals (26) or insect vectors (87).

We found a statistically-significant correlation between recent antibiotic treatment and resistance of faecal bacteria only in the case of gentamicin, and not for sulfa-drugs, trimethoprim, doxycycline, ceftiofur or penicillin, despite the fact that previous literature has found good correlations between antibiotic use and AMR in farm animals (83, 88). An increase in levels of AMR in faecal microbiota of antibiotic-treated horses has been reported previously for *E*.*coli* (83, 85, 89–91), enterococci (89), and gram-negative anaerobes (88), and one study has also reported a lack of resistance in equine *E*.*coli* from wild horses (92). While direct comparisons to these studies are difficult due to methodological differences, the overall trends appear similar. The variables determining the relationship between specific antibiotic treatments and resultant resistances deserve more attention in future studies, and it would be interesting to follow up on some of the unexpected findings from our study such as the apparently increased abundance of some resistance phenotypes (e.g. Sm^R^) in the wild horse cohort compared to the farm and vet cohorts.

Our study appears to be the first to take a ‘taxonomically untargeted’ approach to address questions about AMR and integrons in the equine microflora, using media (R2A) and methods (30°C incubation for 5 days) that allowed the growth of a broad range of aerobic bacteria. This alternative approach allowed us to make the novel observation that many integron-containing Actinobacteria are present in the equine faecal microflora; these would have been overlooked if the samples had been plated on e.g. MacConkey agar. Class 1 integrons have not been seen before in *Glutamibacter*, and only reported once previously in each of the other actinobacterial genera studied here (*Rhodococcus, Arthrobacter, Microbacterium, Micrococcus, Kocuria*) (21, 29, 93). Our study is the first to provide sequence data and genomic context for C1I’s in most of these genera. Class 1 integrons are clearly more common in Actinobacteria than was previously recognised, and furthermore, these occur in the gut of healthy animals in the absence of apparent antibiotic selection. We found evidence for the spread of identical C1I sequences in Actinobacteria between different horses and cohorts; this appears to be due to movement of MGE’s between different bacterial hosts and not just proliferation of clonal bacteria that already carry the C1I’s, but further molecular epidemiology work is needed to confirm this.

A dramatic taxonomic shift was seen in the integron-containing culture collection moving from the farm cohort to the vet cohort, with the latter dominated by *Enterobacteriaceae* and especially *Escherichia spp*. It is worth noting that this trend occurred despite the use of general-purpose isolation media that did not directly select for this species; this emphasises that a continuing focus on *E*.*coli* as a major reservoir of ARGs in equines and other animals is justified. The collection of *Escherichia* strains displayed extensive phenotypic and genotypic diversity, with 41 distinct resistance profiles, 61 distinct BOX-PCR patterns, 18 different resistance genes, and 7 plasmid types detected. Evidence of C1I mobility between *Escherichia* strains and between animals was obtained within the collection, with a single identical 4539 bp contig (*intI1-dfrA17-aadA5-qacEΔ-sul1*) detected in 10 independent isolates from 4 different horses. As was seen in the Actinobacteria, this spread of C1Is appears to be due to movement of MGEs between different bacterial hosts rather than clonal expansion of a single *Escherichia* strain.

Distinct trends were seen in the kinds of MGEs associated with C1Is in the Actinobacteria vs. Proteobacteria isolates. In the Actinobacteria, IS*6100* was the prime mover of C1Is and their associated ARGs, and putative C1I-containing IS*6100-*based composite transposons were found in all these isolates except *Micrococcus*. Intriguingly, these putative transposons were not identical, with several different junction sequences seen between the C1I and the IS element. This could be due to different initial mobilization events, or later internal deletions; in either case, IS*6100* appears to be a very active MGE in these lineages. Although we did not establish the full genomic context of the C1I’s in most of the Actinobacterial isolates, in the case of *Kocuria*, the C1I was located on a novel plasmid with backbone features (RepA, ParA, VirB4/D4/D2) that were very divergent from any previously characterised plasmids; the *Kocuria* contig was also unique in that the complete IS6100-based transposon was recovered, with flanking direct repeats providing clear evidence for its transposition. The *Micrococcus* contig was also unique, firstly because a newly-identified IS element, IS*Mcte1*, was responsible for mobilising the C1I, secondly because the transposon was chromosomal rather than plasmid-borne, and finally because the whole 7 kb CI1-containing mobilised region was near-identical to sequences seen previously in gram negative bacteria, notably In0 from *Pseudomonas* plasmid pVS1, but also >200 other GenBank entries from various enteric bacteria; this provides strong evidence for gene flow from gram negatives to gram positives, resulting in ‘old’ integrons and transposons being mobilised into new contexts.

The MGEs associated with C1Is in our gram negative isolates were more diverse than in the gram positives, but a common theme was the presence of IS*26* in the majority of cases (20/25 contigs). Small IS*26*-based composite transposons (approx. 4 kb) appeared to have mobilised the C1I’s in many of the *Escherichia* strains from the vet cohort horses, and this IS was also present in longer contigs from *Klebsiella, Enterobacter* and *Acinetobacter*. These findings are consistent with the known importance of IS*26* in mobilising ARGs in other contexts (94, 95). Other IS elements seen in the gram negative isolates were IS*1*, IS*1326*, IS*Pa10*, IS*Aba1*, ISCR2, IS*UnCu1*; these are also all known from previous studies to be associated with C1Is and ARGs (67, 96–99). We also found many transposons in our gram negative equine isolates that are known to associate with C1I and ARGs in other contexts (Tn*21*, Tn*402*, Tn*1721*, Tn*7*, Tn*512*), and identified new transposons, notably the putative copper resistance element Tn*7519* and the putative formaldehyde resistance element Tn*7489*. Homologs of Tn*7489* are widely distributed in Enterobactericeae; we propose that this is due to selection presure arising from the use of aldehydes as biocides in diverse medical, agricultural, and industrial contexts. We discovered a new relationship between transposons and integrons in the *Pantoea* contig, ie. mobilisation of the C1I by Tn512 using the IRi site of the integron as one end for transposition.

Our study, like many others, raises questions about why class 1 integrons and resistance genes persist in bacteria in the absence of obvious selection pressures. The most notable example in this study was the *Pantoea* plasmid from a wild horse in a national park that coded for resistances to many heavy metals and formaldehyde in addition to antibiotics, even though none of these toxic agents would be predicted to be present at significant levels in this horse’s environment. Possible explanations include the presence of naturally occurring antibiotics in soil (4), possible alternative benefits from resistance genes (e.g. *sul* and *dfr* genes may assist folate biosynthesis), selection effects at subinhibitory levels (100), co-selection via other useful genes on the same plasmid (101), or the presence of addiction modules like toxin/antitoxin systems (101). Further studies combining molecular ecological and evolutionary approaches with both pure cultures and complex communities are needed to better understand the factors that lead to the persistence of resistance genes in animals and in the environment.

## MATERIALS AND METHODS

Faeces were collected from horses between April 2018 and October 2019 with the approval of the University of Sydney Animal Ethics Committee (project # 2019/1549). Faecal samples were collected from eleven horses at each of three different locations in the state of New South Wales; 1. At a farm in Kembla Grange, where the horses were healthy and had not received antibiotics within six months, 2. At Yarrangobilly in Kosciusko National Park, where the wild horses were of unknown health condition but had never received antibiotic therapy, 3. At Camden Veterinary School, where all horses sampled were being treated for illness, and had recently received antibiotics. Fresh faeces were collected within five minutes of defecation, and sterile tongue-depressor sticks used to remove material from the centre of the faecal deposit to avoid contamination with surrounding soil. Specimens were transferred to sterile Falcon tubes and kept on ice until returned to the laboratory (within 24 hours).

Tenfold dilutions of faecal samples were prepared in sterile water and spread-plated onto R2A agar containing cycloheximide (100 μg/ml) and one of eight different antibiotics, as follows (concentrations in μg/ml); ampicillin (Ap, 100), chloramphenicol (Cm, 25), gentamicin (Gm,10), Kanamycin (Km, 50), rifampicin (Rf, 10), sulfamethoxazole (Sx, 100), streptomycin (Sm, 100), or tetracycline (Tc, 10). Control plates containing cycloheximide only were used to estimate the total number of culturable aerobic heterotrophic bacteria. Plates were incubated for 5 days at 30°C before counting colonies and screening for C1I.

Colonies arising on the antibiotic agars (9000 in total) were picked for further analysis, with these chosen approximately equally among the 3 cohorts and 8 antibiotics. Colonies were duplicated into two 96 well polypropylene plates, the first containing 200 μl phosphate-buffered saline (PBS), and the second containing 200 μl 0.2 M NaOH. The latter plate was heated at 95°C for 15 min to lyse the cells, then 10 μl of lysates from 8 wells were combined, and 1 μl of the mixed lysate used as a template for PCR using primers HS915 and HS916 (42), specific to the *intI1* gene. Any *intI1*^*+*^ isolates were recovered from the PBS cell suspension, restreaked to ensure purity, then stored in 20% glycerol at -80°C for later analysis.

The initial collection of 365 C1I^+^ isolates was dereplicated using BOXA1R PCR (43) before further phenotypic and genotypic analysis. Resistance profiles were determined by a disc diffusion method (44), using 19 antimicrobial discs (Oxoid, details in footnotes of Table 1). The published method was modified to use R2A agar as the base medium, and a 3-day incubation to allow the slower-growing isolates to yield a result. Isolates were identified by Sanger sequencing (Australian Genome Research Facility, Westmead, NSW, Australia) of PCR-amplified partial 16S rRNA genes (primers 27F and 519R (45), followed by blastn analysis against the NCBI nucleotide database (46). Gene cassette arrays were amplified using primers HS458 and HS459 (47), with a 5 min extension time sufficient to recover arrays of up to 10 kb, and also Sanger-sequenced. Draft whole genomes were obtained from a subset of isolates using Illumina sequencing by Microbial Genome Sequencing Center, Pittsburgh, PA, USA (approx. 2.1 million reads per isolate) and the UTS Core Sequencing Facility (University of Technology, Sydney, NSW, Australia (approx. 2.7 million reads per isolate); DNA for this analysis was prepared via a beadbeating and silica binding method (48).

Programs on the Galaxy bioinformatics server (release v 21.09) were used for initial processing and analysis of raw genomics data; these included FASTQC v0.73 for quality assessment and quality control (49), CutAdapt v4.0 (50) and Trimmomatic v0.36.6 (51) for removal of adapters, SPAdes v3.15.4 (52) for assembly of reads, and Prokka v1.14.6 (53) for initial genome annotation. PCR-amplified cassette arrays and genome contigs containing C1Is were further analysed using Abricate (54) and the Comprehensive Antibiotic Resistance Database (55) to identify ARGs, PlasmidFinder (56) to detect plasmid replication functions and assign incompatibility groups, and the INTEGRALL database (21) to identify integron gene cassettes. Contigs were then uploaded into SnapGene (57) for more detailed curation including BLASTp of individual ORFs, comparison to the NCBI Conserved Domain Database, and manual identification of *attC* sites, inverted repeats, direct repeats and iterons. SPSS Statistics v26.0.0.0 (IBM) was used for statistical analyses, with either ‘non-parametric legacy test on 2 independent samples’ or ‘1-tailed exact significance’ tests used dependent on the data set. Graphs were produced with GraphPad v8.4.3 (Prism).

Annotated integron-containing genome contigs from the equine faecal isolates have been deposited in GenBank with accession numbers OP296261-OP296273.

## ACKNOWLEDGEMENTS

Scott Mitchell was supported by an Australian Government Research Training Program Scholarship. This project was also supported by the University of Sydney CDIP Industry and Community Engagement fund grant (#CT21044), which included a financial contribution from Quantal Bioscience Pty Ltd. We thank Valentina Vitale, Linda Lord, and Darryl Nelson for assisting with access to horses.

